# *Arid1a* loss potentiates pancreatic β-cell regeneration through activation of EGF signaling

**DOI:** 10.1101/2020.02.10.942615

**Authors:** Cemre Celen, Jen-Chieh Chuang, Shunli Shen, Jordan E. Otto, Clayton K. Collings, Xin Luo, Lin Li, Yunguan Wang, Zixi Wang, Yuemeng Jia, Xuxu Sun, Ibrahim Nassour, Jiyoung Park, Alexandra Ghaben, Tao Wang, Sam C. Wang, Philipp E. Scherer, Cigall Kadoch, Hao Zhu

## Abstract

The dynamic regulation of β-cell abundance is poorly understood. Since chromatin remodeling plays critical roles in liver regeneration, these mechanisms could be generally important for regeneration in other tissues. Here we show that the ARID1A mammalian SWI/SNF complex subunit is a critical regulator of β-cell regeneration. *Arid1a* is highly expressed in quiescent β-cells but is physiologically suppressed when β-cells proliferate during pregnancy or after pancreas resection. Whole-body *Arid1a* knockout mice were protected against streptozotocin induced diabetes. Cell-type and temporally specific genetic dissection showed that β-cell specific *Arid1a* deletion could potentiate β-cell regeneration in multiple contexts. Transcriptomic and epigenomic profiling of mutant islets revealed increased Neuregulin-ERBB-NR4A signaling. Functionally, *ERBB3* overexpression in β-cells was sufficient to protect against diabetes, and chemical inhibition of ERBB or NR4A was able to block increased regeneration associated with *Arid1a* loss. mSWI/SNF complex activity is a barrier to β-cell regeneration in physiologic and disease states.

## Introduction

Understanding how to recover endogenous β-cells in disease might ultimately depend on understanding how β-cell abundance is controlled in the first place. The molecular circuitry that dictates total β-cell mass during development and dynamic physiologic situations such as pregnancy is poorly understood. It is also unclear if the same mechanisms regulate β-cell expansion after injuries and during disease processes. One unexploited therapeutic strategy is to expand endogenous β-cells in type 1 and 2 diabetes, settings in which normal regenerative capacity cannot fully compensate for β-cell loss. Given the importance of dedifferentiation and proliferation in profoundly regenerative organisms such as zebrafish and planaria (Jopling et al., 2011), it stands to reason that facilitating chromatin state changes that occur physiologically could also increase self-renewal capacity in mammalian tissues. We hypothesized that the mammalian SWI/SNF (mSWI/SNF) ATP-dependent chromatin remodeling complex, known to promote terminal differentiation (Han et al., 2019; Hota et al., 2019; Sun et al., 2016b; Yu et al., 2013; Zhang et al., 2019, 2016) and also known to limit regeneration after liver injuries (Sun et al., 2016b), might operate as a general repressor of regeneration in tissues other than the liver. Given key similarities between β-cell and hepatocyte self-renewal, we reasoned that mSWI/SNF might also suppress regenerative capacity in β-cells, a centrally important cell type in metabolic disease.

mSWI/SNF are large 10-15 component complexes containing a core ATPase (BRG1 or BRM, also known as SMARCA4 and SMARCA2), plus non-catalytic subunits that influence targeting and complex activities (Wu et al., 2009) such as the mutually exclusive ARID1A (BAF250A) and ARID1B (BAF250B) subunits, which together define the canonical BAF family of mSWI/SNF complexes. The loss of ARID1A alters the assembly of the ATPase module (Mashtalir et al., 2018), influencing canonical BAF complex targeting and DNA accessibility, resulting in global changes in gene regulation (Chandler et al., 2013; Kelso et al., 2017; Mathur et al., 2017; Nakayama et al., 2017; Pan et al., 2019; Sun et al., 2018). How the molecular consequences of mSWI/SNF perturbation relate to physiologic phenotypes and disease outcomes is an active area of exploration.

Intriguingly, ARID1A and other mSWI/SNF components are downregulated during islet expansion, findings shared with other regenerative tissues such as the liver (Sun et al., 2016b). We demonstrated that this downregulation is functionally important by using spatially and temporally specific conditional knockout mouse models subjected to multiple types of β-cell injuries. These findings support the idea that physiological events that occur during regeneration can be further amplified to accelerate tissue healing. Molecular dissection of the events occurring downstream of ARID1A loss showed that NRG-ERBB-NR4A signaling activities were increased. Interestingly, the ERBB protein family (EGFR/ERBB1, ERBB2, ERBB3, and ERBB4) of transmembrane receptor tyrosine kinases (RTKs) have been implicated in diabetes (Miettinen et al., 2006; Oh et al., 2011; Song et al., 2016). Remarkably, SNPs in the human *ERBB3* locus are among the strongest signals in T1D GWAS but how these SNPs affect ERBB signalling and β-cell biology were previously unknown (Bradfield et al., 2011; Nikitin et al., 2010; Sun et al., 2016a). Here, we show that ARID1A-containing SWI/SNF complexes have a major and unexpected role in β-cell regeneration, operating in large part through pathways that were previously implicated by GWAS studies in diabetogenesis. This expands our emerging understanding of mSWI/SNF as a central regulator of tissue regeneration, and underscores the need to understand chromatin remodeling mechanisms that may represent therapeutic targets in regeneration.

## Results

### *Arid1a* expression is suppressed during physiologic β-cell expansion

ARID1A-containing SWI/SNF chromatin remodeling complexes (or canonical BAF complexes) drive terminal differentiation and block proliferation by regulating chromatin accessibility at loci targeted by lineage specific transcription factors (Alver et al., 2017; Spaeth et al., 2019; Vierbuchen et al., 2017). In the absence of ARID1A, increased liver regeneration and accelerated wound healing were observed (Sun et al., 2016b). Although the primary mechanism for the homeostatic maintenance of adult β-cells is self-duplication, majority of adult pancreatic β-cells are still mostly in post-mitotic state (Dor et al., 2004; Meier et al., 2008). We wondered if the inhibition of SWI/SNF activity could increase their proliferative potential. Genes that encode components of the SWI/SNF complex are expressed at high levels in the mouse β-cell line MIN6 and primary pancreatic islets compared to the mouse α-cell line ATC1 and primary hepatocytes (**Figure 1A**). In particular, *Arid1a* is highly expressed in quiescent β-cells within the islet (**Figure 1B**). Next, we examined *Arid1a* expression in islets under conditions that demand β-cell expansion, such as pregnancy and 50% partial pancreatectomy (PPx) (Ackermann Misfeldt et al., 2008; Rieck and Kaestner, 2010). *Arid1a* mRNA was lower in maternal islets during pregnancy-associated β-cell expansion, and protein levels declined until the time of birth (**Figure 1C,D**). Similarly, ARID1A protein levels were reduced during regeneration induced by PPx (**Figure 1E**).

**Figure 1.**
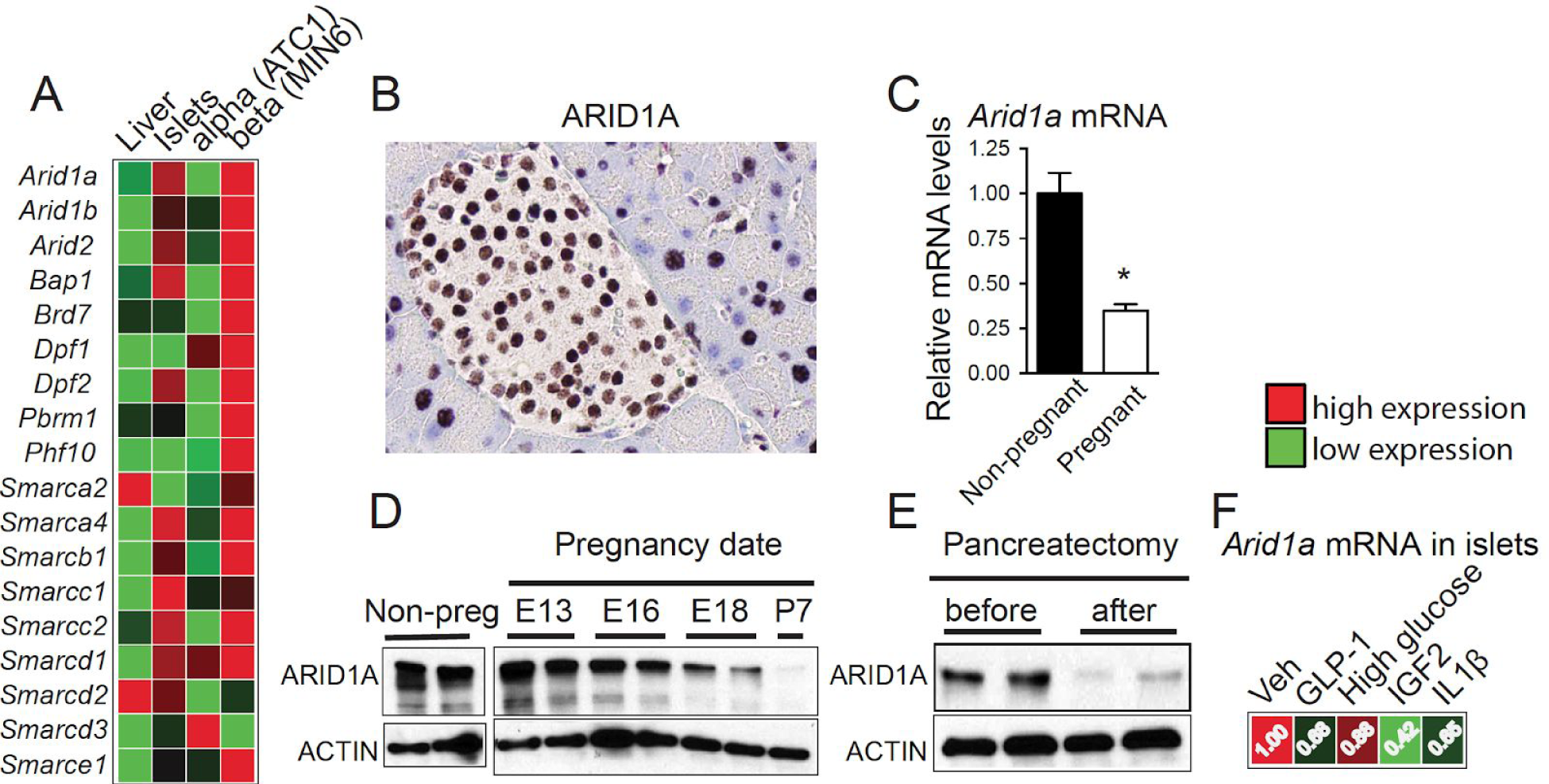
*Arid1a* expression is suppressed during physiologic β-cell expansion. **A.** mRNA expression heat map of SWI/SNF components in mouse liver, mouse islet, alpha cell line (ATC1) and beta cell line (MIN6) using qPCR. **B.** ARID1A immunostaining in the adult mouse pancreas. An islet is shown. **C.** qPCR showing *Arid1a* mRNA levels in islets isolated from non-pregnant and pregnant females at gestational day 16 (n = 3 mice per group). **D.** Western blot showing ARID1A levels in the islets of non-pregnant females and at different time points during gestation. **E.** Western blot showing ARID1A levels in the islets before and 5 days after 50% pancreatectomy (PPX). **F.** qPCR showing the reduction in *Arid1a* levels in isolated islets after 6 hours of treatment in culture with the following stimuli: GLP-1 (100 nM), high glucose (25 mM), IGF2 (200nM) and IL1β (1 ng/ml).

We asked if signals known to elicit β-cell hyperplasia can influence the levels of *Arid1a* mRNA. We isolated wild-type (WT) mouse islets and treated with glucagon-like peptide-1 (GLP-1), high glucose, insulin-like growth factor-2 (IGF2), and interleukin-1β (IL-1β), all factors known to drive β-cell proliferation (Alismail and Jin, 2014; De León et al., 2003; Gaddy et al., 2010; Hajmrle et al., 2016; Porat et al., 2011). Each factor suppressed *Arid1a* mRNA levels in islets, providing a potential molecular rationale for how the suppression of *Arid1a* during β-cell expansion is linked to upstream signals (**Figure 1F**). These results indicated dynamic regulation of ARID1A and SWI/SNF complex components during β-cell expansion and regeneration, suggesting functional roles during these processes.

### Whole-body *Arid1a* deletion protected against type I diabetes

To determine if ARID1A restrains β-cell mass in adults, we inducibly deleted *Arid1a* in a whole body “*Arid1a* KO” mouse strain. Global deletion was induced with tamoxifen in adult *Ubiquitin-CreER; Arid1a^Fl/Fl^* (*Ubc-CreER; Arid1a^Fl/Fl^*) mice (**Figure 2A**). Almost complete absence of the flox band and protein levels were observed (**Figure 2B-D**). Whole body *Arid1a* KO mice did not show overt signs of disease and appeared to be healthy. At baseline, whole body glucose tolerance was unchanged, indicating that ARID1A is not required for the regulation of glucose homeostasis or uptake under non-injury, basal conditions (**Figure 2E**). To determine if *Arid1a* KO mice were more able to cope with β-cell destruction in a type 1 diabetes (T1D) model, we subjected mice to streptozotocin (STZ), a chemical that ablates insulin-producing β-cells (**Figure 2A**). After STZ, *Arid1a* KO mice were almost completely protected against the development of diabetes as measured by fed state glucose levels (**Figure 2F**) and glucose tolerance testing (GTT) (**Figure 2G**). This left the possibility of either insulin secretion or insulin sensitivity changes, particularly because *Arid1a* was deleted in all tissues. Mice subjected to STZ did not show changes in insulin sensitivity based on insulin tolerance testing (ITT), suggesting no substantial metabolic impact of *Arid1a* loss on peripheral tissues, despite efficient deletion in those tissues (**Figure 2H**). However, *Arid1a* KO mice produced higher levels of insulin after glucose administration (**Figure 2I**), suggesting a β-cell mediated mechanism of protection against or response to STZ-induced T1D.

**Figure 2.**
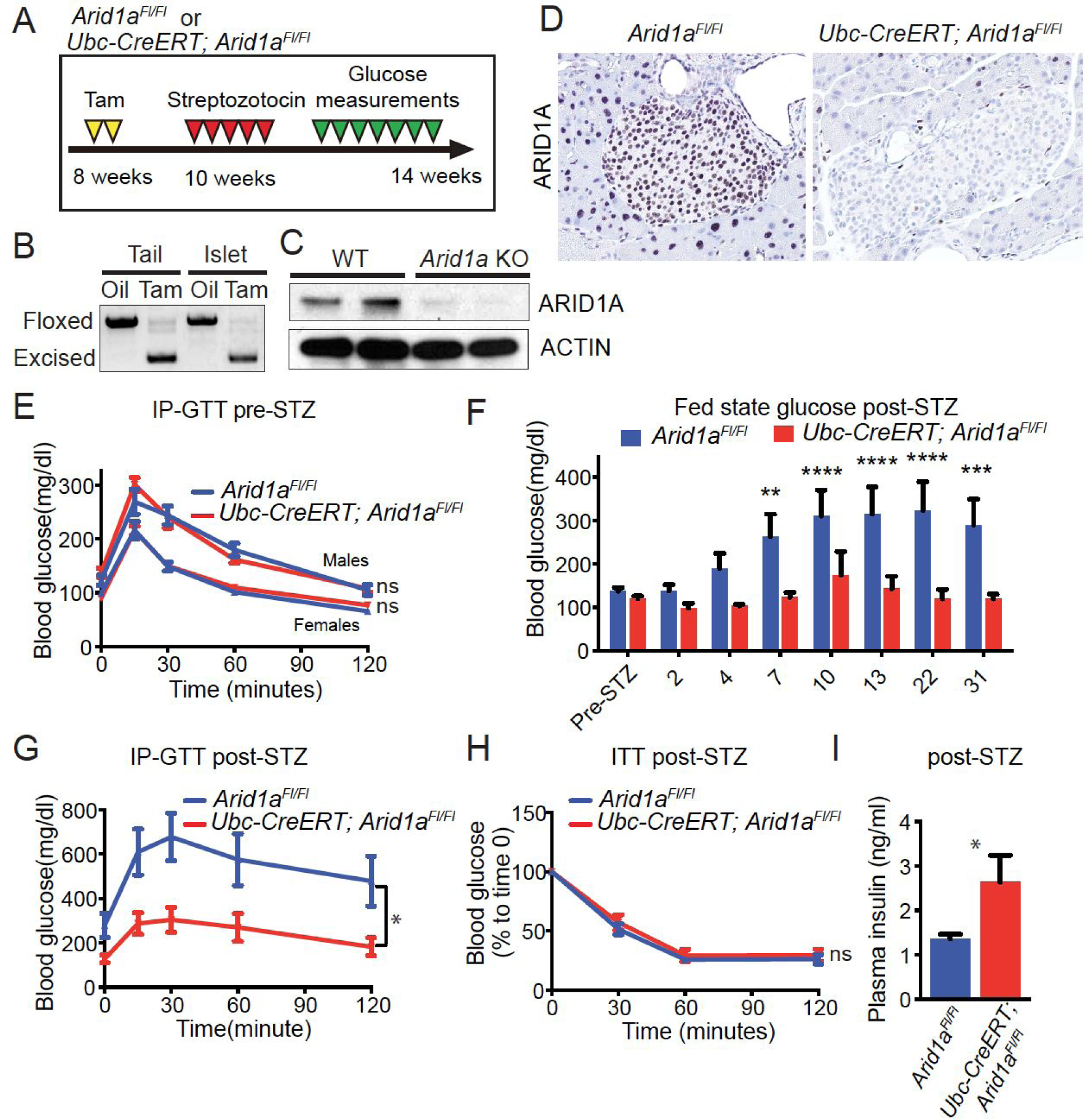
Whole body *Arid1a* deletion protects against STZ-induced T1D. **A.** Schema for STZ-induced diabetes studies in *Arid1a^Fl/Fl^* control and *Ubc-CreER; Arid1a^Fl/Fl^* whole body KO mice. Two weeks after administration of 20 mg/kg tamoxifen oral gavage x 2 days, diabetes was induced by injecting 50 mg/kg STZ IP for 5 consecutive days. **B.** Genotyping of tail and islets to assess the recombination at the *Arid1a^Fl/Fl^* locus. **C.** Western blot assessing the reduction in ARID1A protein levels. **D.** Immunostaining of ARID1A in the adult pancreas of *Ubc-CreER; Arid1a^Fl/Fl^* mice. **E.** IP glucose tolerance test at baseline (Females: n=5 for *Arid1a^Fl/Fl^* and n=4 for *Ubc-CreER; Arid1a^Fl/Fl^*, Males: n=3 for *Arid1a^Fl/Fl^* and n=8 for *Ubc-CreER; Arid1a^Fl/Fl^*). **F.** After STZ, fed state blood glucose was measured following STZ (n=11 for *Arid1a^Fl/Fl^* and n=9 for *Ubc-CreER; Arid1a^Fl/Fl^*). **G.** IP glucose tolerance test performed 2 weeks post-STZ. **H.** Insulin tolerance test performed 2 weeks post-STZ. **I.** Plasma insulin levels measured by ELISA 2 weeks post-STZ.

### *Arid1a* deficiency leads to a β-cell-mediated anti-diabetic phenotype

Given that the whole body KO mice have lost *Arid1a* in multiple cell types, it was possible that non-β-cell autonomous mechanisms might have been at play. In WT mice, ARID1A is expressed in most acinar, all duct, and all islet cells. To rule out potential paracrine or endocrine effects, we used *Ptf1a-Cre; Arid1a^Fl/Fl^* mice to induce deletion in non-β-cells in the pancreas. *Ptf1a* is a transcription factor that is expressed in all cell types in the pancreas starting around E9.5 (Kawaguchi et al., 2002). In *Ptf1a-Cre; Arid1a^Fl/Fl^* mice, ARID1A was lost in all acinar and most duct cells but ARID1A protein was retained in the islets, as shown recently (Wang et al., 2019a, 2019b). GTT did not reveal any differences in glucose clearance efficiency between WT, *Ptf1a-Cre; Arid1a^+/Fl^*, or *Ptf1a-Cre; Arid1a^Fl/Fl^* mice (**Figure S1A**). Following STZ treatment (**Figure S1B**), there were also no differences in blood glucose measurements over the course of 1 month (**Figure S1C)**, ruling out a potential contribution from acinar or duct cells to the phenotypes observed in whole-body KO mice.

Next, we generated a temporally regulated, β-cell specific model of *Arid1a* loss. This model allowed us to induce β-cell specific deletion of *Arid1a* after β-cell injuries, thus giving us the ability to answer questions specifically about β-cell regeneration, rather than cell survival in the presence of injuries. We employed a *MIP (Mouse Insulin Promoter)-rtTA; TRE (Tetracycline Responsive Element promoter)-Cre* transgenic system to allow such genetic control (**Figure 3A**) (Kusminski et al., 2016). *MIP-rtTA; TRE-Cre; Arid1a^Fl/Fl^ (Arid1a* βKO) mice and littermate controls without *Cre* and/or *rtTA* were used to model cell-type and temporally specific *Arid1a* deletion. Notably, some *Arid1a* floxed band was still present in islets, potentially due to α, δ and γ cell retention of the *Arid1a* floxed allele (**Figure 3B**). Despite the absence of ARID1A nuclear staining (**Figure 3C**), islet cell morphology was normal. Two weeks post-deletion, the ability to clear glucose after GTT challenge was unaltered (**Figure 3D**), suggesting that there was no change in β-cell function at baseline. Next, both groups of mice were given five doses of STZ (50mg/kg per day x 5 days), then fed doxycycline (1mg/mL) water to conditionally induce *Arid1a* deletion in the β-cells of the experimental group. β-cell specific deletion of *Arid1a* after STZ administration prevented the rise of blood glucose to pathological levels (**Figure 3E**), and reduced the profound weight loss associated with T1D (**Figure 3F**). In the setting of STZ, *Arid1a* βKO showed higher insulin levels than control mice. Not surprisingly, insulin levels were still lower in the STZ treated *Arid1a* βKO group than uninjured WT mice since STZ was still able to ablate a subset of β-cells (**Figure 3G**). These data showed that *Arid1a* deficiency in β-cells was sufficient to preserve insulin production in response to STZ-induced T1D.

**Figure 3.**
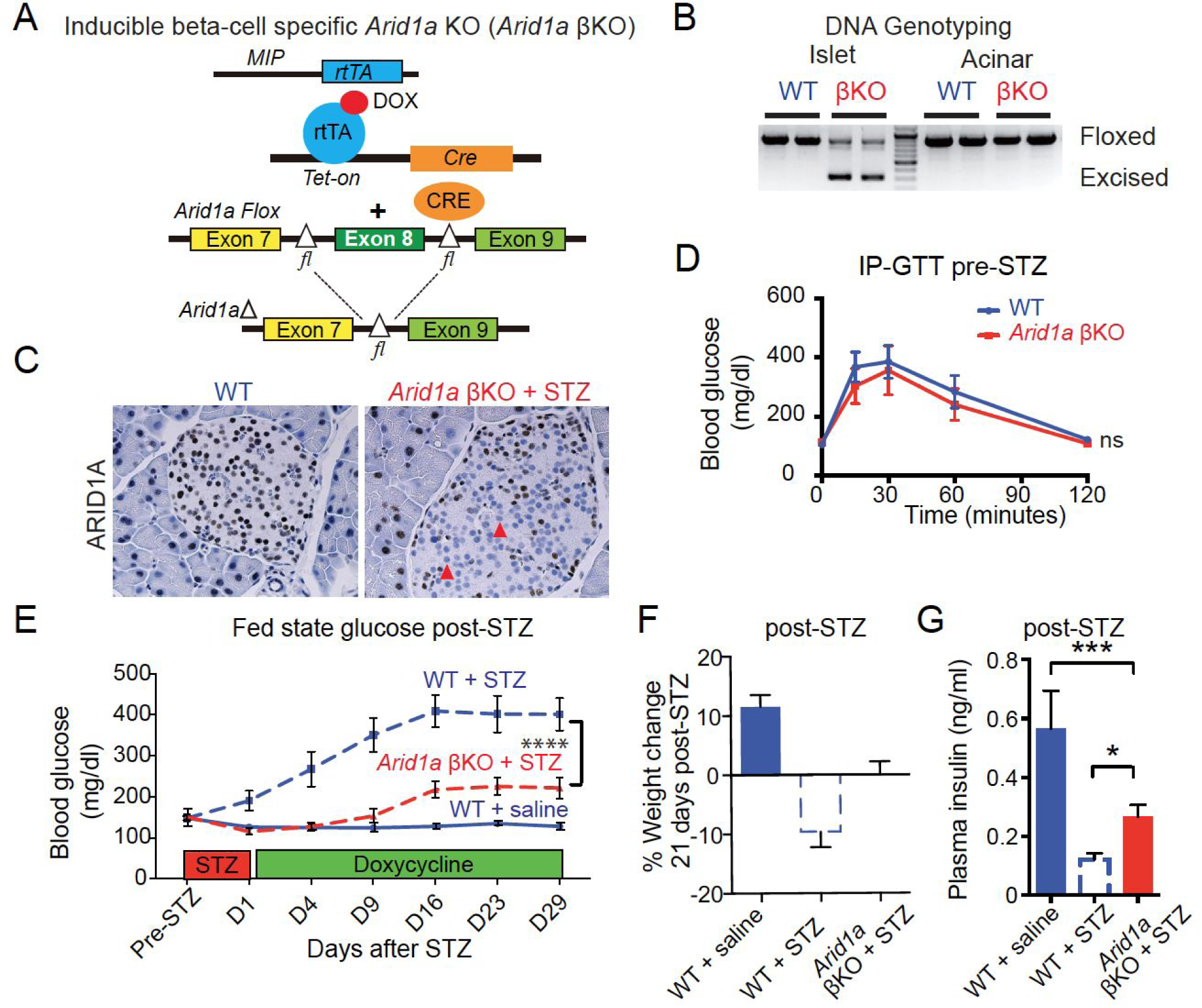
*Arid1a* deficiency leads to a β-cell-mediated anti-diabetic phenotype. **A.** Dox-inducible KO of *Arid1a* in mouse β-cells (*Arid1a* βKO). rtTA is expressed under the control of the mouse insulin promoter (*MIP*). In the presence of doxycycline (Dox), rtTA activates the transcription of the *TRE-Cre* transgene. CRE in turn converts the floxed *Arid1a* alleles to knockout alleles. Dox (1mg/mL) was added to the drinking water to conditionally induce *Arid1a* deletion. **B.** Genotyping of islets and acinar cells to assess the recombination at the *Arid1a^Fl/Fl^* locus. Partial excision is expected in islets since cell types other than β cells are present. **C.** Immunostaining of ARID1A in WT and *Arid1a* βKO pancreata. **D.** IP glucose tolerance test 2 weeks post-dox. **E.** Fed state blood glucose after STZ. 5 doses of 50 mg/kg STZ were injected before dox-induced β-cell deletion of *Arid1a*. After the last dose of STZ, dox (1mg/mL) was provided in drinking water (n=4 mice for WT + saline, n=12 mice for WT + STZ, n=10 mice for *Arid1a* βKO + STZ). **F.** % body weight change 21 days post-STZ. **G.** Plasma insulin levels measured by ELISA 30 days post-STZ.

### *Arid1a* loss results in increased β-cell survival and proliferation

We examined islets before and after STZ injury. Insulin staining in WT and *Arid1a* βKO islets at baseline showed no differences. Thirty days after STZ, *Arid1a* βKO mice had more β-cells than did WT mice (**Figure 4A**). In addition, WT mice showed considerable α-cell expansion as marked by glucagon staining (**Figure 4B**). Because resistance to diabetes could have been mediated by either a failure to lose β-cells or an increased ability to regenerate β-cells, we examined proliferation in the insulin expressing β-cell compartment. *Arid1a* deficient β-cells had greater numbers of Ki-67 positive cells. In the WT setting, a majority of the proliferating cells were positioned in the islet periphery, which is the non-insulin expressing compartment (**Figure 4C,D**). We then wondered whether the observed increase in proliferation after STZ had any effect on islet number or area. Although we did not detect any increase in the number of islets per unit of pancreas area (**Figure 4E**), there was a trend towards increased individual islet area (p=0.06) (**Figure 4F**) in βKO mice 3 weeks after STZ.

**Figure 4.**
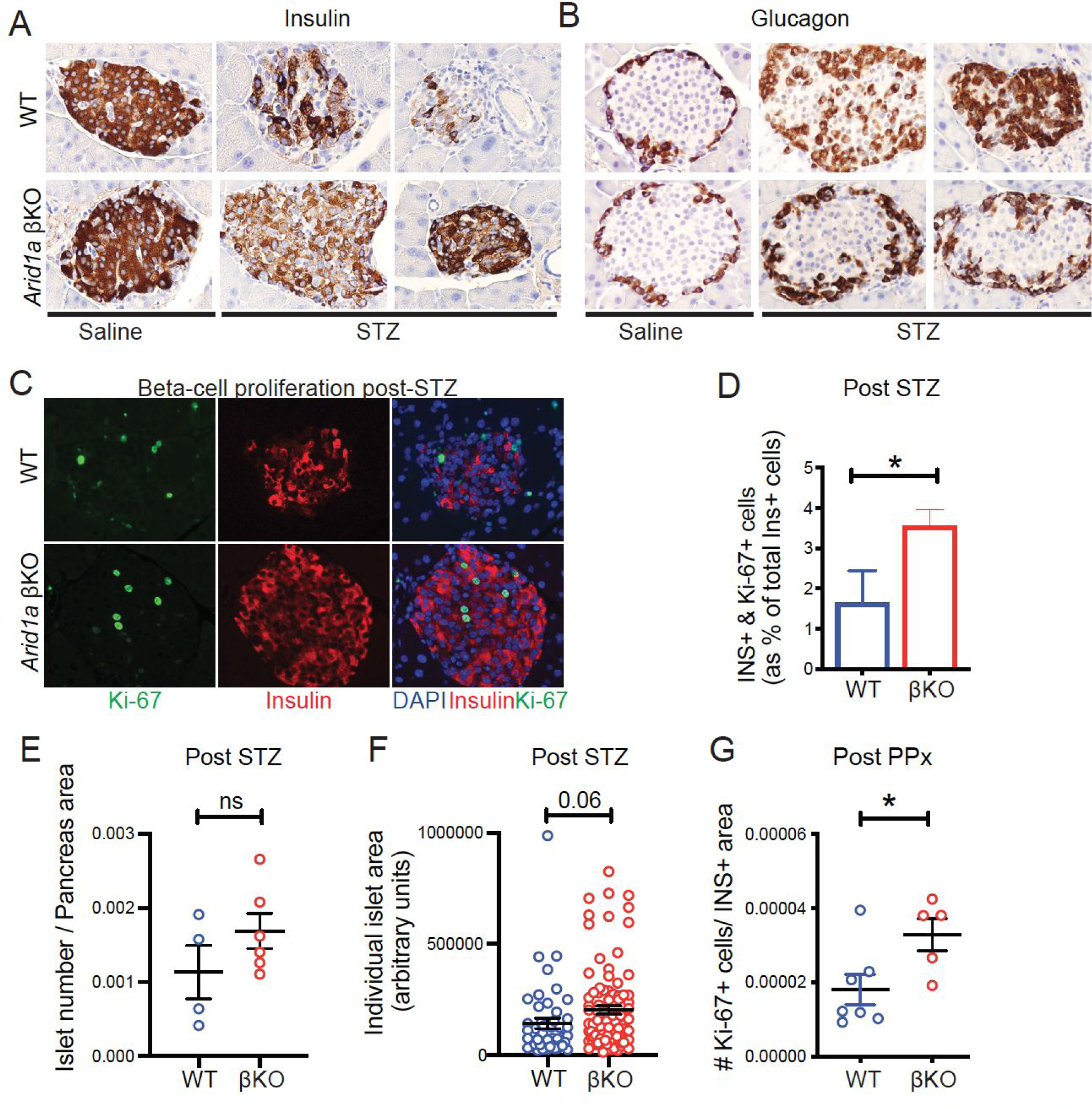
*Arid1a* loss results in increased β-cell survival and proliferation. **A.** Insulin staining of WT and *Arid1a* βKO pancreata treated with saline or STZ. Samples were collected 1 month post-STZ. **B.** Glucagon staining of WT and *Arid1a* βKO pancreata treated with saline or STZ. Samples were collected 1 month post-STZ. **C.** Representative Ki-67 staining (green) of WT and *Arid1a* βKO pancreata post-STZ. Insulin is used as a β-cell marker (red). Samples were collected 1 month post-STZ. **D.** Quantification of the percentage of insulin+ & Ki-67+ cells to all insulin+ cells (total of 9 islets from 1 WT and total of 25 islets from 3 *Arid1a* βKO mice) **E.** The number of islets per pancreas section area in WT and *Arid1a* βKO mice 3 weeks post-STZ. Each dot represents the total number of islets per section area in each mouse (n = 4 WT mice for WT and n = 6 *Arid1a* βKO mice). **F.** Individual islet area (n = 4 WT mice and 6 *Arid1a* βKO mice, at least 10 islets were measured for each mouse). **G.** Ki-67+ cell number/insulin+ area in islets from regenerated pancreata 6 days after PPx (n = 7 WT and 5 *Arid1a* βKO mice, values from ∼10 islets per mice were calculated and averaged to get a single data point in the graph).

Given the remote possibility that *Arid1a* deficient β-cells could be protected from T1D due to a lack of STZ induced destruction rather than increased regeneration, we also performed PPx, a surgical assay that does not rely on the ability of cells to metabolize chemical toxins such as STZ. PPx leads to β-cell proliferation that compensates for overall islet loss. Six days after resection, the islets from regenerated pancreata of βKO mice showed a significant increase in β-cell proliferation as measured by Ki-67 positive cell number within the insulin expressing compartment (**Figure 4G**). In summary, *Arid1a* loss in β-cells did not constitutively enforce islet overgrowth in the absence of injury but increased proliferation after chemical and surgical injuries.

### Phenotypes associated with *Arid1a* loss are dependent on EGF/Neuregulin hyperactivation

To probe the transcriptional programs associated with *Arid1a* deletion, we performed RNA-seq on islets from control and KO mice, before and after pancreatectomy. Of 2796 differentially-expressed genes, 1586 were up and 1210 were downregulated (**Figure 5A**). Gene Set Enrichment Analysis (GSEA) performed on differentially expressed genes showed that genes involved in the neuregulin (NRG) and epidermal growth factor (EGF) pathways were collectively overproduced in KO islets (**Figure 5B & Figure 5A right inset**). EGF and NRG are the ligands that activate ERBB family of RTKs. Binding of the EGF family growth factors leads to dimerization and phosphorylation of RTKs which leads to the activation of ERK1/2 and phosphatidylinositol 3-kinase (PI3K) signaling pathways (Yarden and Sliwkowski, 2001).

**Figure 5.**
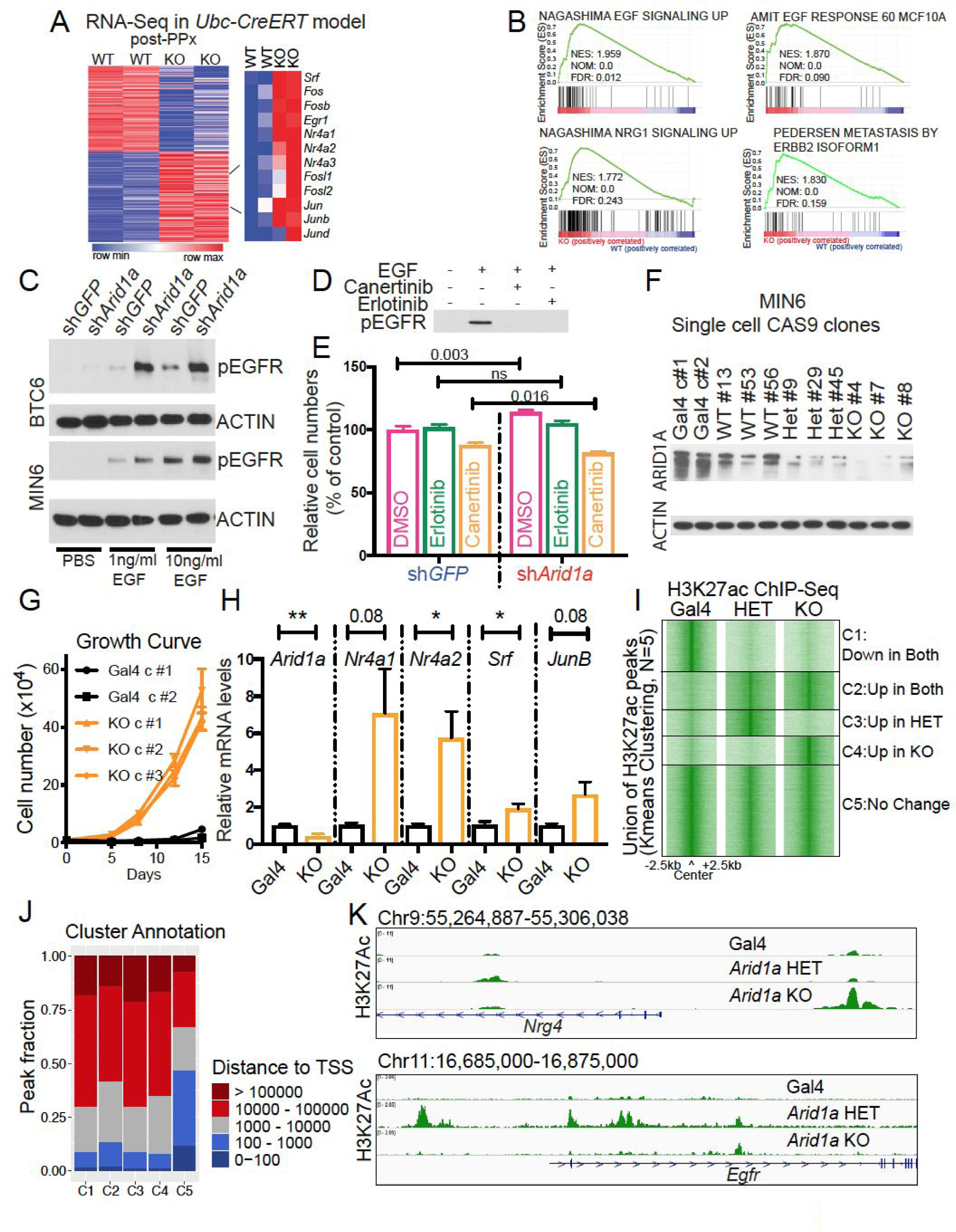
*Arid1a* deficient β-cells have a dependence on increased EGF/Neuregulin signaling. **A.** Heatmap of differentially expressed genes in WT and KO islets from *Arid1a^Fl/Fl^* and *Ubc-CreER; Arid1a^Fl/Fl^* mice, isolated after PPx and detected by RNA-Seq (left). Heatmap showing a subset of overexpressed genes in KO islets (right; n = 2 and 2 mice). **B.** GSEA shows that EGF and NRG1 response genes are upregulated in KO islets. The nominal enrichment score (NES), nominal p-value, and false discovery rate (FDR) q-value are shown within each GSEA plot. **C.** p-EGFR western blots in MIN6 and BTC6 treated with different doses of EGF. **D.** p-EGFR western blots in WT MIN6 cells treated with EGF in the presence or absence of erlotinib and canertinib. **E.** Relative cell numbers for control *shGFP* and *shArid1a* MIN6 cells in the presence of canertinib and erlotinib. Shown as % of control cells (*shGFP* group treated with DMSO). **F.** Western blot showing ARID1A levels in MIN6 clones. MIN6 cells were transduced with lenti-*CAS9*-blasticidin and either non-targeting lenti-sgRNA (*Gal4*)-puromycin or lenti-sgRNA (*Arid1a*)-puromycin. **G.** MIN6 clone growth over 15 days measured by cell counting. **H.** mRNA expression of selected genes in *Gal4* control and *Arid1a* KO MIN6 clones as measured by qPCR. **I.** K-mean clustering of H3K27Ac ChIP-Seq peaks in non-targeting *Gal4*, *Arid1a* heterozygous and *Arid1a* KO MIN6 clones. **J.** Annotation of clusters defined by H3K27Ac ChIP-Seq sites by distance to TSS. **K.** Sample tracks for H3K27Ac ChIP-Seq in control, *Arid1a* heterozygous and *Arid1a* KO MIN6 single cell clones at the *Nrg4* and *Egfr* loci.

To functionally interrogate the transcriptomic findings, we performed shRNA knockdown of *Arid1a* in BTC6 and MIN6 immortalized β-cell lines to determine if *Arid1a* loss might cause increased dependency on NRG/EGF signaling. *Arid1a* knockdown in BTC6 and MIN6 caused increased phospho-EGFR (p-EGFR) after EGF ligand exposure, indicating potentiation of the pathway (**Figure 5C**). In addition, EGFR/ERBB inhibition with the small molecule inhibitors erlotinib and canertinib were able to ablate p-EGFR in the presence of EGF (**Figure 5D**). Similar proliferative phenotypes were confirmed with *Arid1a* shRNA in MIN6 cells and these small molecule inhibitors selectively abrogated the hyperproliferation associated with *Arid1a* knockdown (**Figure 5E**).

To corroborate the shRNA results using another *in vitro* model, we generated *Arid1a* deficient MIN6 β-cells using CRISPR. Multiple single cell MIN6 clones for each genotype were expanded and confirmed for *Arid1a* loss (**Figure 5F**). Similar to β-cells from mouse islets, *Arid1a* KO MIN6 clones grew more rapidly than Gal4 targeted control clones (**Figure 5G**). Genes identified in the RNA-seq experiment were also upregulated in KO vs. control islets (*Nr4a1, Nr4a2, Srf, JunB*; see **Figure 5H**). Collectively, these data show that *Arid1a* loss in β-cells causes a preferential dependency on ERBB signalling.

To determine if differentially expressed genes from the RNA-Seq analysis also showed changes in epigenetic marks, we performed H3K27 acetylation (H3K27ac) ChIP-seq. Sites marked by H3K27ac were clustered into groups that increased or decreased in H3K27ac in response to *Arid1a* deletion (**Figure 5I**). Interestingly, many sites increased in H3K27ac abundance in the *Arid1a* heterozygous and homozygous clones (clusters 3 and 4). The clusters containing sites that changed in H3K27ac occupancy upon *Arid1a* loss were largely composed of sites at distal regions (**Figure 5J)**, which is consistent with the disruption of canonical BAF complexes (Kelso et al., 2017; Mathur et al., 2017; Vierbuchen et al., 2017). Sites that increased in H3K27ac upon *Arid1a* loss included *Nrg4* and *Egfr*, consistent with transcriptional hyperactivation in the *Arid1a*-deficient setting (**Figure 5K)**. GO biological processes associated with increased acetylation of H3K27 in the *Arid1a* knockout populations in clusters 3 and 4 contain terms involving glucocorticoid biosynthesis and hormone metabolism, and the associated mouse phenotypes included abnormal insulin secretion and pancreas secretion (**Figure S2A** and **Figure S2B**). These observations are consistent with differential regulation of pathways affecting glucose metabolism and insulin secretion in KO mouse β-cells.

To challenge the idea that *Arid1a* deletion causes more proliferation through the preferential promotion of ERBB signaling as opposed to a more generalized activation of multiple mitogenic signal pathways, we performed a small molecule screen to identify pathways that *Arid1a* KO cells are more dependent on for growth and survival. We screened 300 kinase inhibitors on previously generated immortalized H2.35 cells isogenic for *Arid1a* deletion (**Figure S3A**). Interestingly, ERBB family inhibitors were significantly enriched among treatments that induced the most prominent reductions in cell viability (P-value = 5.8e-6). There were 3 ERBB family inhibitors (Afatinib, WZ8040, canertinib) among the top 10 kinase inhibitors associated with the highest reduction of growth/viability in *Arid1a* KO cells (**Figure S3B**). Together with the islet transcriptome data, these observations established a strong functional connection between ARID1A and ERBB.

### *ERBB3* overexpression independently increases β-cell regeneration

In order to perform gain-of-function studies for ERBB signalling, we first asked which *ERBB* receptor loci ARID1A might be acting through. ChIP-seq data on human liver cell line HepG2 showed binding of ARID1A and core SWI/SNF component SNF5 on the promoters of *ERBB2* and *ERBB3* (**Figure 6A**) (Raab et al., 2015). In addition, multiple human GWAS studies have shown strong associations between a SNP (rs2292239) in *ERBB3* and T1D (Barrett et al., 2009; Kaur et al., 2016; Sun et al., 2016a; Todd et al., 2007). Based on these observations, we reasoned that ARID1A could have a genetic interaction with *ERBB3*. Because it is still unknown whether or not an increase or decrease of *ERBB3* in β-cells or other tissues is protective in human T1D, we generated a β-cell specific human *ERBB3* overexpression mouse model (*MIP-rtTA; TRE-ERBB3*) (**Figure 6B**). Strikingly, inducible overexpression of *ERBB3* alone was sufficient to mediate resistance to STZ-induced T1D (**Figure 6C**). This is the first functional confirmation of an effect for *ERBB3* on a T1D model.

**Figure 6.**
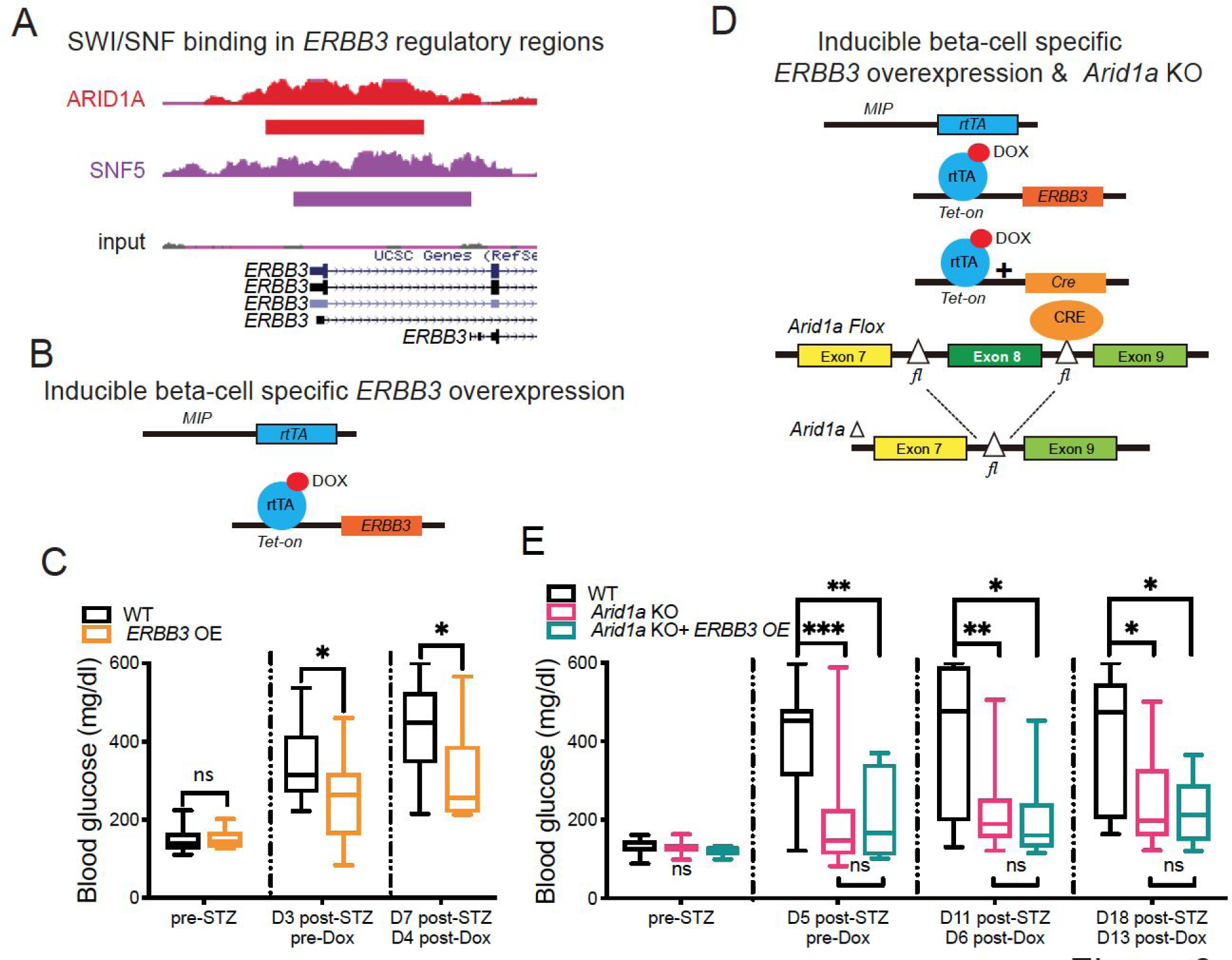
*ERBB3* overexpression in β-cells is protective against T1D after STZ. **A.** Binding of ARID1A and SNF5, a core SWI/SNF subunit, to the *ERBB3* locus in HepG2 cells (ChIP-seq data from (Raab et al., 2015)). **B.** Schema of dox-inducible β-cell specific *ERBB3* overexpression mice: *MIP-rtTA*; *TRE-ERBB3*. **C.** Fed state blood glucose measurements in control and *ERBB3* overexpressing mice post-STZ. **D.** Schema of dox-inducible β-cell specific *ERBB3* overexpression and *Arid1a* βKO mice: *MIP-rtTA*; *TRE-ERBB3; TRE-Cre; Arid1a^fl/fl^*. **E.** Fed state blood glucose measurements in WT, *Arid1a* βKO and *Arid1a* βKO + *ERBB3* OE mice post-STZ.

To determine if there is an epistatic relationship between *ERBB3* and *Arid1a*, we genetically manipulated both genes in vivo. We hypothesized that if the two genes act via distinct pathways, *ERBB3* overexpression in an *Arid1a* deficient background would have a synergistic effect (**Figure 6D**), whereas if both genes are within the same pathway, then there would be little to no additive effect. While mice with either β-cell specific *ERBB3* overexpression or *Arid1a* loss showed protection against STZ-induced diabetes, simultaneous perturbations did not synergize to increase the protective effect (**Figure 6E**). The lack of a synergistic effect suggests that either these genes are acting in the same pathway, or that the effects are individually saturated. These results for the first time validate the human GWAS signals for *ERBB3* in an animal model and support the hypothesis that *Arid1a* loss is operating through increases in ERBB signaling.

### Pan-ERBB and NR4A1 inhibition ablated the *Arid1a* KO phenotype in vivo

We sought to determine whether pro-proliferative effects of *Arid1a* loss were mediated through ERBB activation in vivo as well as in vitro. We first tested the in vivo relevance of these results by examining whole body *Ubc-CreER; Arid1a* WT and KO mice described earlier. After STZ mediated islet ablation, mice were given daily doses of canertinib, an intervention that abolished the anti-diabetic effects of *Arid1a* loss (**Figure 7A**, compare to **Figure 2F** which did not include canertinib). We reasoned that ERBB inhibition in the STZ model could have been influenced by cell death in addition to proliferation, so we also performed the PPx assay to assess β-cell proliferation after surgical injury. Here, β-cell specific *Arid1a* deletion was induced one week before PPx and drug treatments were started one day before and continued until 6 days after PPx, which is the apex for β-cell proliferation (Peshavaria et al., 2006). The increase in Ki-67/insulin double positive β-cells became more pronounced in the regenerating pancreas (**Figure 7B, top row & 7C**). Again, canertinib abolished the pro-proliferative effect seen in KO islets (**Figure 7B, middle row & 7C**). These results showed that ERBB signaling in part mediates the β-cell regenerating effects of *Arid1a* deficiency in vivo.

**Figure 7.**
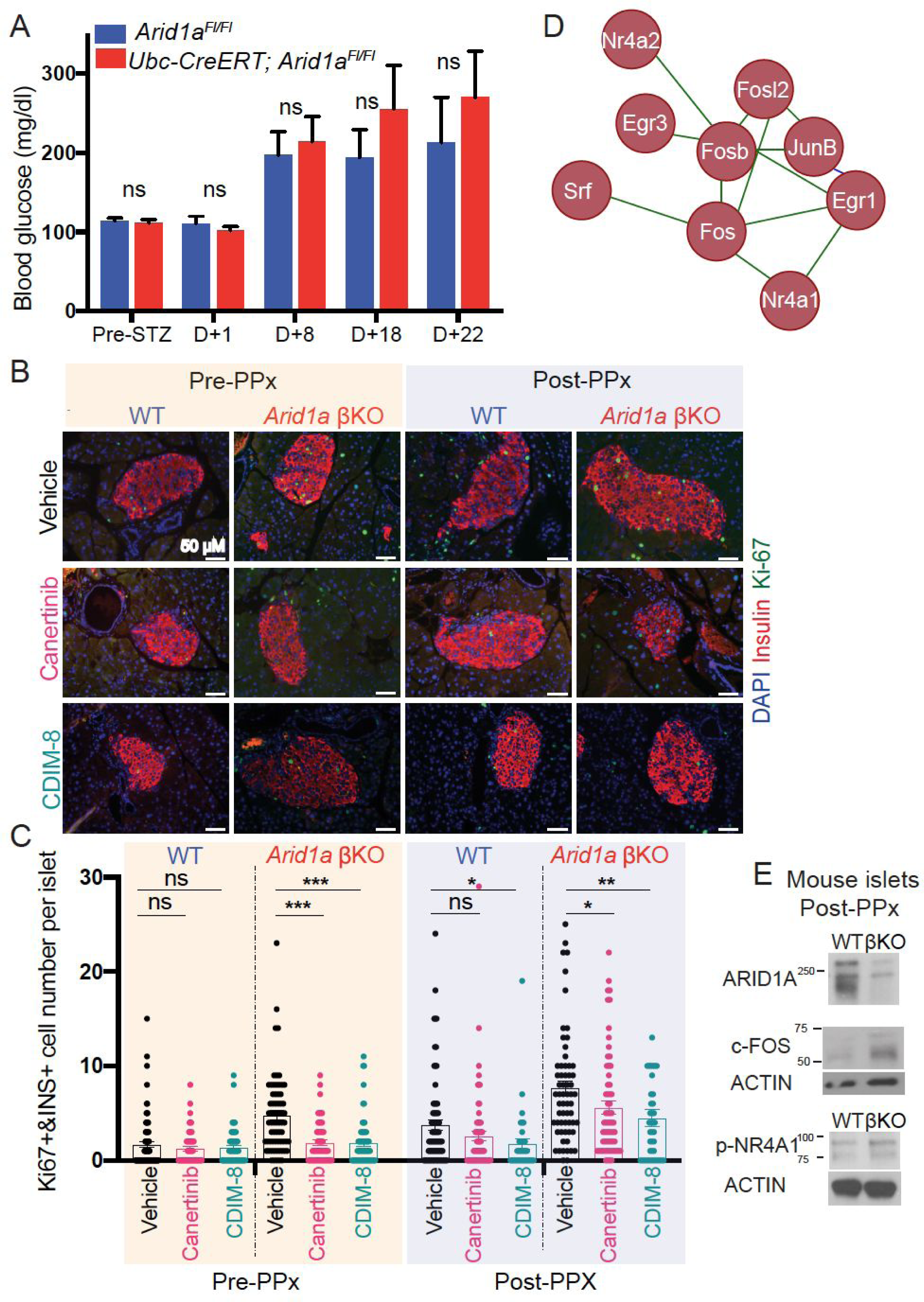
Pan-ERBB and NR4A1 inhibition abrogated the *Arid1a* KO phenotype in vivo. **A.** Fed state blood glucose measurements following STZ in WT and whole body *Arid1a* KO mice. 20mg/kg canertinib was administered daily through oral gavage. Compare to Fig. 2F, which did not include canertinib. **B.** Immunofluorescence for DAPI (blue), insulin (red), and Ki-67 (green) in WT and *Arid1a* βKO islets before or after PPx in the presence of vehicle, canertinib, or CDIM-8 treatment. **C.** Quantification of Ki-67 and insulin double positive cell number per islet. Each dot represents an islet. Between 37-75 islets per genotype were counted from 5 to 8 mice per genotype. **D.** The STRING database predicts an important protein-protein interaction network from differentially overexpressed genes found in RNA-seq data. **E.** Western blot analysis of c-FOS and p-NR4A1 in mouse islets isolated 6 day post-PPx.

Because ARID1A likely influences a large network of genes that regulate differentiation and regenerative capacity, we next asked if other genes downstream of ERBB signaling exert the effects of *Arid1a* loss. To further examine important functional networks resulting *Arid1a* KO phenotype, a protein-protein interaction network of overexpressed genes from the RNA-seq data was created using the STRING database (Szklarczyk et al., 2015). This interaction network showed that the SRF/JUN/FOS/EGR1 transcription factors and the NR4A family of nuclear receptor transcription factors were tightly connected and clustered together (**Figure 7D**). Indeed, these genes also came up under the GSEA datasets that were classified as “EGF/NRG1 signaling up” (**Figure 5A**) suggesting that they are downstream of ERBB signaling. The overexpression of these transcription factors were previously established as one of the hallmarks of immature, proliferative β-cells (Zeng et al., 2017). We validated higher protein levels of c-FOS and p-NR4A1 in KO islets (**Figure 7E**). We then asked if ARID1A relays its effects in part through NR4A1 in vivo by treating mice that had undergone PPx with an NR4A1 antagonist (C-DIM8), in a similar fashion as the canertinib experiments. C-DIM8 also abolished the increased proliferation normally seen in *Arid1a* KO β-cells (**Figure 7B, bottom row & 7C**), suggesting that increased NR4A1 activity is additionally required for the phenotypic effects of *Arid1a* loss.

## Discussion

Preserving or replenishing endogenous pancreatic β-cells is a major challenge in diabetes. To ultimately meet these goals, there is a need to understand the factors limiting β-cell proliferation. Although engineering strategies such as inducible pluripotent stem cell to β-cell conversion or the transdifferentiation of other cell types to β-cells are emerging (Aguayo-Mazzucato and Bonner-Weir, 2018), targeting endogenous regeneration of β-cells has potential advantages since this occurs in development, pregnancy, and obesity to meet increased physiological demands for insulin production. Here, we have identified *Arid1a* as a factor whose suppression is required for gestational β-cell expansion and is sufficient for β-cell replenishment after multiple pancreatic injuries. Previous studies have shown that BRG1 and BRM-containing SWI/SNF complexes have context-dependent roles in modulating PDX1 activity in β-cell development and function (McKenna et al., 2015; Spaeth et al., 2019). Wei *et. al.* reported that Vitamin D ligand causes the Vitamin D receptor to interact with an active BRD7/pBAF rather than an inactive BRD9/ncBAF complex, which reduces β-cell failure and slows diabetes progression (Wei et al., 2018). Our study is the first to highlight *Arid1a* and SWI/SNF as an important epigenetic node that regulates the regenerative capacity of β-cells.

By functionally validating EGF/NRG1 signaling as a critical effector of ARID1A suppression, we connected two previously unrelated, but prominent regeneration networks. EGF signaling is known to be important for liver, heart, and pancreas regeneration (D’Uva et al., 2015; Gemberling et al., 2015; Song et al., 2016; Sun et al., 2016b). GWAS studies implicated *ERBB3* as one of the most significant T1D gene loci, but it is not clear if *ERBB3* expression is positively or negatively influencing T1D risk, and if *ERBB3* is acting in β-cells, immune cells, or peripheral tissues (Barrett et al., 2009; Kaur et al., 2016; Sun et al., 2016a; Todd et al., 2007). For the first time, we determined that *ERBB3* overexpression in β-cells is sufficient to protect against diabetes. The fact that overexpression did not add to the protective phenotype of *Arid1a* deletion is consistent with the idea that ARID1A loss acts through ERBB signaling.

Mechanistically, our results implicate a network of EGF/NRG1 signaling genes. These include *Nr4a* nuclear receptor genes and a set of nutrient-responsive immediate early genes (*JunB*, *Fos*, *Egr1*) and their upstream activator *Srf* (Mina et al., 2015). Interestingly, EGR binding sites represented the most dynamic chromatin regions during whole body regeneration of *Hofstenia miamia*, an acoel worm that is capable of whole body regeneration. In these worms, EGR is essential for regeneration through its activities as a pioneer factor that directly regulated wound-induced genes including the EGFR ligands *nrg-1* and *nrg-2* (Gehrke et al., 2019). In addition, Zeng et al. recently used single cell transcriptomics to categorize β-cells from less to most mature. They showed that *Srf*/*JunB*/*Fos*/*Egr1* were among the most downregulated genes during postnatal β-cell maturation and comprise a signature of less mature, proliferative β-cells. Importantly, overexpression of upstream activator *Srf* in islets did not impair insulin secretion (Zeng et al., 2017). Our data suggest that ARID1A containing complexes may negatively regulate this larger network of pro-regeneration genes. Importantly, β-cells without ARID1A are poised for proliferation while still able to maintain insulin production.

Our study highlights targets that could be investigated for their therapeutic potential in future studies. In contrast to the majority of nuclear receptors, the NR4A family of nuclear receptors do not have an identified physiological ligand, therefore their activity is mainly regulated at the level of protein expression and post-translational modifications (Safe et al., 2016). However, there is a agonist, Cytosporone-B, that can selectively increase the transcriptional activity of *Nr4a*1 (Zhan et al., 2008). Whether this agonist could be utilized to improve the regenerative potential of β-cells is an area of interest. A more direct way to tackle this problem would be to use specific inhibitors targeting ARID1A or BAF complexes. There is a major interest in the field to develop small molecules to manipulate activities of SWI/SNF complex, and if successful, it will be interesting to determine if such inhibitors might be effective in diabetes.

## Acknowledgements

We would like to thank Michael Kalwat and Melanie H. Cobb for sharing MIN6 cell line. Histology Pathology Core (John Shelton), CRI Sequencing Core (Jian Xu, Xin Liu, Yoon Jung Kim) and CRI Mouse Genome Engineering Core (Yu Zhang) contributed to this project. We would like to thank Eric Olson, Jiang Wu, and Jian Xu for their valuable feedback during this project. C.C. and X.S. were supported by The Hamon Center for Regenerative Science and Medicine (CRSM) Trainee Fellowships. T.W. is supported by a R03ES026397-01 and CPRIT (RP150596). H.Z. was supported by the Pollack Foundation, an NIH/NIDDK R01 grant (DK111588), a Burroughs Wellcome Career Award for Medical Scientists, a CPRIT Scholar Award (R1209).

## Author Contributions

C.C. and J.C. designed and performed the experiments and wrote the paper. S.S. maintained the mouse colony and performed surgical procedures, islet isolations and data analysis. J.E.O and C.K.C performed H3K27Ac ChIP-Seq experiments. X.L. and Y.W. performed and T.W. supervised the bioinformatics analysis. L.L. generated MIN6 single cell clones and performed experiments. Z.W., X.S., I.N., J.P., A.G., P.E.S, S.C.W generated and analyzed mouse models. S.C.W. and C.K. edited the manuscript. H.Z. conceived and supervised the project and wrote the manuscript.

## STAR Methods

### LEAD CONTACT AND MATERIALS AVAILABILITY

Further information and material requests should be directed to Dr. Hao Zhu.

### EXPERIMENTAL MODEL AND SUBJECT DETAILS

#### Mice

All experiments on mice were approved by and handled in accordance with the guidelines of the Institutional Animal Care and Use Committee at UTSW. All experiments were performed in in age and gender controlled fashion. Male mice were used for STZ experiments. The transgenic mouse lines used are as follows and described before: *Arid1a* floxed mice (JAX stock #027717) (Gao et al., 2008)*, Ptf1a-Cre* (Kawaguchi et al., 2002), *MIP-rtTA* (Kusminski et al., 2016)*, TRE-Cre* (JAX 006234) (Perl et al., 2002)*. TRE-HER3* mouse line was generated by Jiyoung Park and Alexandra Ghaben in Dr. Scherer’s group and a paper with the detailed characterization of this mouse model is in preparation.

#### Cell Lines

The H2.35 (ATCC® CRL-1995™) and BTC6 cells were obtained from ATCC (ATCC® CRL-11506™) and cultured according to manufacturer’s protocol. MIN6 cell line was obtained from Dr. Melanie Cobb’s lab and cultured in DMEM with 15% Heat Inactivated Fetal Bovine Serum, 1% L-Glutamine, 1% Pen/Strep, 0.0005% beta mercaptoethanol.

#### Mouse Islet Culture

Islets from adult mice were isolated and recovered overnight in culture medium (RPMI1640 with 10% heat-inactivated FBS, L-glutamine and Pen/Strep) in the incubator and used for experiments.

## METHOD DETAILS

### Mouse pancreatic islet isolation and dispersion of islets

Islet isolation was done as described previously (Zmuda et al., 2011) with minor modifications. Briefly, Liberase TL (Roche, 05401020001) solution was prepared by dissolving 5 mg lyophilized Liberase TL powder in 1 ml sterile water so that the concentration is 5 mg/ml corresponding to 26 Wunsch units. Prior to use, this 1 ml Liberase solution was added to 21.6 ml serum free RPMI to obtain the working solution. Adult mice were perfused with 3 ml Liberase TL working solution through the common bile duct cannulation and inflated pancreas was put into 2 ml Liberase TL solution sitting on ice in 50 ml falcon tube and incubated at 37 °C water bath for 10-16 minutes by shaking and reaction was stopped with the addition of RPMI containing serum. Islets were dissociated from the exocrine tissues by shaking vigorously several times followed by Histopaque 1077 (Sigma-Aldrich, 10771) density centrifugation (900g, 20 minutes, acceleration:2, deceleration: 0). Purified islets were collected from the interface and washed. Intact islets were hand-picked under the dissection microscope.

### Islet mitogen experiments

Islets from male CD1 mice were isolated and recovered overnight in culture medium (RPMI1640 with 10% heat-inactivated FBS, L-glutamine and Pen/Strep) in the incubator prior to various treatments. After recovering, islets were treated with either vehicle (culture medium with 11.1 mM glucose), high glucose (culture medium with 25 mM glucose), or with the addition of IGF2 (200nM), GLP-1 (100nM), IL1β (1 ng/ml) for 6 hours and RNA was harvested for qPCR analysis.

### Kinase inhibitor screen

SelleckChem customized library (Z49127) was a collection of 304 kinases inhibitors. See the supplementary table for a full list of inhibitors with additional details about their targets. Isogenic WT and Arid1a KO H2.35 transformed mouse liver hepatocyte cells were used in this kinase screen and treated with kinase inhibitors. To determine if EGFR family inhibitors is among the treatment conditions that induced the highest decrease in cell viability, we performed a hypergeometric test to examine the enrichment of treatments involving these inhibitors. Briefly, we ranked all treatment conditions in descending order according to observed cell viability loss, then, we counted the number of EGFR family inhibitor treatments among the top 10% conditions. The possibility of observing at least N EGFR family inhibitor treatments by chance is calculated and reported as the p-value of EGFR enrichment.

### Generation of CAS9 single cell clones

Mouse *Arid1a* gRNA (GCTGCTGCTGATACGAAGGTTGG) was cloned into LentiGuide-puro plasmid (Addgene #53963). LentiCas9-Blast plasmid was purchased from Addgene (Addgene #53962). Active lentivirus was prepared in 293T cells in 10-cm dish. The day after seeding cells, each dish was transfected with pVSV-G, pLenti-gag pol, LentiCas9-puro-sG*Arid1a* or LentiCas9-Blast plasmid by Lipofectamine 3000 (Life Technologies # L3000015). Virus containing medium was collected at 60h after transfection. For creating the Cas9 stable expressing cell line, cells were infected with Cas9 lentivirus, followed by blasticidin selection at 2ug/ml for 4 days. Then Cas9 expressing cells were infected with Arid1a gRNA lentivirus. Three days after the infection, we selected cells for 3 days with puromycin at 2ug/ml. Then, 50 to 200 cells were plated in 15cm dishes for single clone selection. Single clones were picked when they grew big enough and verified the genotype and Arid1a expression by PCR and Western blot.

### In vivo drugs

Dox water (1g/L) was used to induce conditional deletion. Tamoxifen was dissolved (Sigma-Aldrich, T5648) in corn oil at a concentration of 20 mg/ml by sonication. 500uL of 20mg/mL Tamoxifen by oral gavage for two consecutive days. 20 mg/kg canertinib (LC Laboratories NC0704940) was administered daily through oral gavage for one month in STZ experiments. Either 20 mg/kg canertinib or 20 mg/kg NR4A1 antagonist C-DIM8-DIM-C-pPhOH (Axon Medchem # Axon 2827) was administered daily through gavage starting a day before PPx until sacrificing mice day 7 post-PPx.

### Streptozotocin (STZ) injury

Mice were fasted for 4-6 hours prior to STZ injection. STZ (Sigma-Aldrich S0130) was dissolved in sodium-citrate (Sigma-Aldrich S4641) solution to a final concentration of 10 mg/ml freshly 10-15 min before the injection. Sodium-citrate solution was prepared by dissolving 1.47 gram of sodium-citrate powder in 50 ml ddH2O, and adjusting the pH to 4.5. Prepared STZ solution was injected via intraperitoneal injection. Different dosage was used for different strain backgrounds as indicated in the figure legends since response to STZ is strain dependent.

### Glucose tolerance test (GTT)

After 16 hours fast, blood glucose was measured using a glucometer from tail tip blood before and multiple times within 2 hours of intraperitoneal injection of 2g/kg D-(+)-Glucose (Sigma Aldrich # G7528).

### Insulin tolerance test (ITT)

After 6h fast, blood glucose was measured before intraperitoneal injection of insulin (0.75 mU/g body wt) and then 30, 60, 90 and 120 min after injection.

### Partial pancreatectomy (PPx)

PPx was performed as described (Martín et al., 1999) except that mice were not fasted overnight.

### RNA Extraction and RT-qPCR

Total RNA from isolated mouse islets were extracted using TRIzol reagent (Invitrogen) or The RNeasy Plus Micro Kit (cat. no. 74034). For RT-qPCR of mRNAs, cDNA synthesis was performed with 1 mg of total RNA using miScript II Reverse Transcription Kit (QIAGEN). See Supplemental Information for primers used in these experiments. Expression was measured using the ΔΔCt method.

### Western blot

Isolated islets were lysed in RIPA buffer (Sigma R0278) supplemented with protease and phosphatase inhibitors. Protein concentration was determined by Pierce™ BCA Protein Assay Kit (Thermo Fisher #23225). Western blots were performed in the standard fashion. The following antibodies were used: ARID1A (Sigma-Aldrich Cat# HPA005456, RRID:AB_1078205), ARID1A (Santa Cruz Biotechnology Cat# sc-32761, RRID:AB_673396), p-NR4A1 (Cell Signaling Technology Cat# 5095, RRID:AB_10695108), phospho-EGFR (Tyr1068) (Cell Signaling Technology Cat# 3777, RRID:AB_2096270), phospho-EGFR (Y1173) (Cell Signaling Technology Cat# 4407, RRID:AB_331795), phospho-AKT (Ser473) (Cell Signaling Technology Cat# 4060, RRID:AB_2315049), phospho-p44/42 (p-ERK) (Cell Signaling Technology Cat# 9101, RRID:AB_331646), p44/42 (ERK) (Cell Signaling Technology Cat# 9107, RRID:AB_10695739), c-FOS (Santa Cruz Biotechnology Cat# sc-166940, RRID:AB_10609634), β-Actin (Cell Signaling Technology Cat# 4970, RRID:AB_2223172), Vinculin (Cell Signaling Technology Cat# 13901, RRID:AB_2728768), anti-rabbit IgG, HRP-linked antibody (Cell Signaling Technology Cat# 7074, RRID:AB_2099233) and anti-mouse IgG, HRP-linked antibody (Cell Signaling Technology Cat# 7076, RRID:AB_330924).

### Histology, immunohistochemistry, and immunofluorescence

Tissue samples were fixed in 4% paraformaldehyde (PFA) and embedded in paraffin. H&E staining was performed by the UTSW Histology Core Facility. Primary antibodies used: anti-rabbit Arid1a (Sigma-Aldrich Cat# HPA005456, RRID:AB_1078205, used for IHC), anti-rabbit Glucagon (Cell Signaling Technology Cat# 2760, RRID:AB_659831, used for IHC), anti-rabbit Insulin (Abcam Cat# ab108326, RRID:AB_10861152, used for IHC), anti-rabbit Ki-67 (Abcam Cat# ab15580, RRID:AB_443209, used for IF), anti-guinea pig Insulin (Abcam Cat# ab7842, RRID:AB_306130, used for IF). Secondary antibodies used: Alexa Fluor 488 goat anti-rabbit IgG (Thermo Fisher; 11008; RRID:AB_10563748) goat polyclonal antibody to anti-Guinea pig IgG-Alexa 568 (Abcam, ab175714). For IHC, detection was performed with the Elite ABC Kit and DAB Substrate (Vector Laboratories) followed by hematoxylin counterstaining (Sigma). VECTASHIELD® Antifade Mounting Media with DAPI counterstain (Vector Labs, H-1200) was used. For islet number and area calculation, H&E sections were imaged using The NanoZoomer 2.0-HT whole slide imaging. Number of islets were counted in each slide. Whole H&E section area and area of each individual islet in that H&E section were measured using NDP.view software. To determine the proliferation, slices of pancreas were costained with insulin, Ki-67 and DAPI and all the islets present in the section were imaged using the same parameters. Channels were merged and images were analyzed in Image J.

### RNA-sequencing

RNA was extracted from islets isolated from 2 *Arid1a^Fl/Fl^ and 2 Ubiquitin-CreER; Arid1a^Fl/Fl^* mice. RNA-seq libraries were prepared with the Ovation RNA-Seq Systems 1-16 (Nugen) and indexed libraries were multiplexed in a single flow cell and underwent 75 base pair single-end sequencing on an Illumina NextSeq500 using the High Output kit v2 (75 cycles) at the UTSW Children’s Research Institute Sequencing Facility.

## QUANTIFICATION AND STATISTICAL ANALYSIS

### RNA-Seq Analysis

RNA-Seq analysis was performed as described before (Celen et al., 2017). Briefly, adaptors and low quality sequences (Phred<20) were removed by trimming raw sequencing reads using galore package (http://www.bioinformatics.babraham.ac.uk/projects/trim_galore/). The sequence reads were aligned to the GRCm38/mm10 with HiSAT2 (Kim et al., 2015). After duplicates removal by SAMtools (Li et al., 2009), read counts were generated for the annotated genes based on GENCODE V20, using featureCounts (Harrow et al., 2012; Liao et al., 2014). Differential gene expression analysis was performed on edgeR, using FDR < 0.05 as cutoff (McCarthy et al., 2012; Robinson et al., 2010). Heatmaps to visualize the data were generated by using GENE-E (Robinson et al., 2010).

### GSEA

GSEA was used to determine significantly enriched gene sets. To perform GSEA analysis using RNA-seq data from WT and *Arid1a* KO islets, raw read counts from each sample were converted to cpm value (count per million) using the cpm function within edgeR R package. GSEA analysis was performed with a pre-ranked gene list by log fold change. GSEA was then performed against hallmark gene sets using default parameters (http://software.broadinstitute.org/gsea/index.jsp).

### ChIP-seq Data Processing and Analysis

Sequencing of ChIP-seq samples was carried out using the Illumina NextSeq technology, and reads were demultiplexed with the bcl2fastq software tool. Read quality was evaluated by FASTQC, and read alignment to the hg19 genome was executed with Bowtie2 v2.29 in the -k 1 reporting mode (Langmead and Salzberg, 2012). Narrow peaks were detected using MACS2 v2.1.1 with a q-value cutoff of 0.001 and input as controls (Zhang et al., 2008). Output BAM files were transformed into BigWig track files using the “callpeak” function of MACS2 v2.1.1 with the “-B --SPMR” option followed by the use of the BEDTools (Quinlan and Hall, 2010) “sort” function and the UCSC utility “bedgraphToBigWig”. BigWig track files were then input in IGV v2.4.3 for visualization. Heat maps were generated using ngsplot v2.61, which was also used to perform K-means clustering (Shen et al., 2014). Cis-regulatory function analysis was carried out using the GREAT online software suite (Argiras et al., 1987).

### Statistical analysis

Variation is indicated using standard error presented as mean ± SEM. Two-tailed Student’s *t*-tests (two-sample equal variance) were used to test the significance of differences between the two groups. Statistical significance is displayed as * (P < 0.05), ** (P < 0.01), *** (P < 0.001), ****(P < 0.0001). Statistical analyses were performed using GraphPad Prism unless otherwise indicated. Mice from multiple litters were used in the experiments. In STZ follow up experiments, mice were occasionally excluded from the analysis due to death.

## DATA AND CODE AVAILABILITY

All sequencing data have been deposited in the GEO with the accession number (pending).

## FIGURES AND LEGENDS

**Figure S1.**
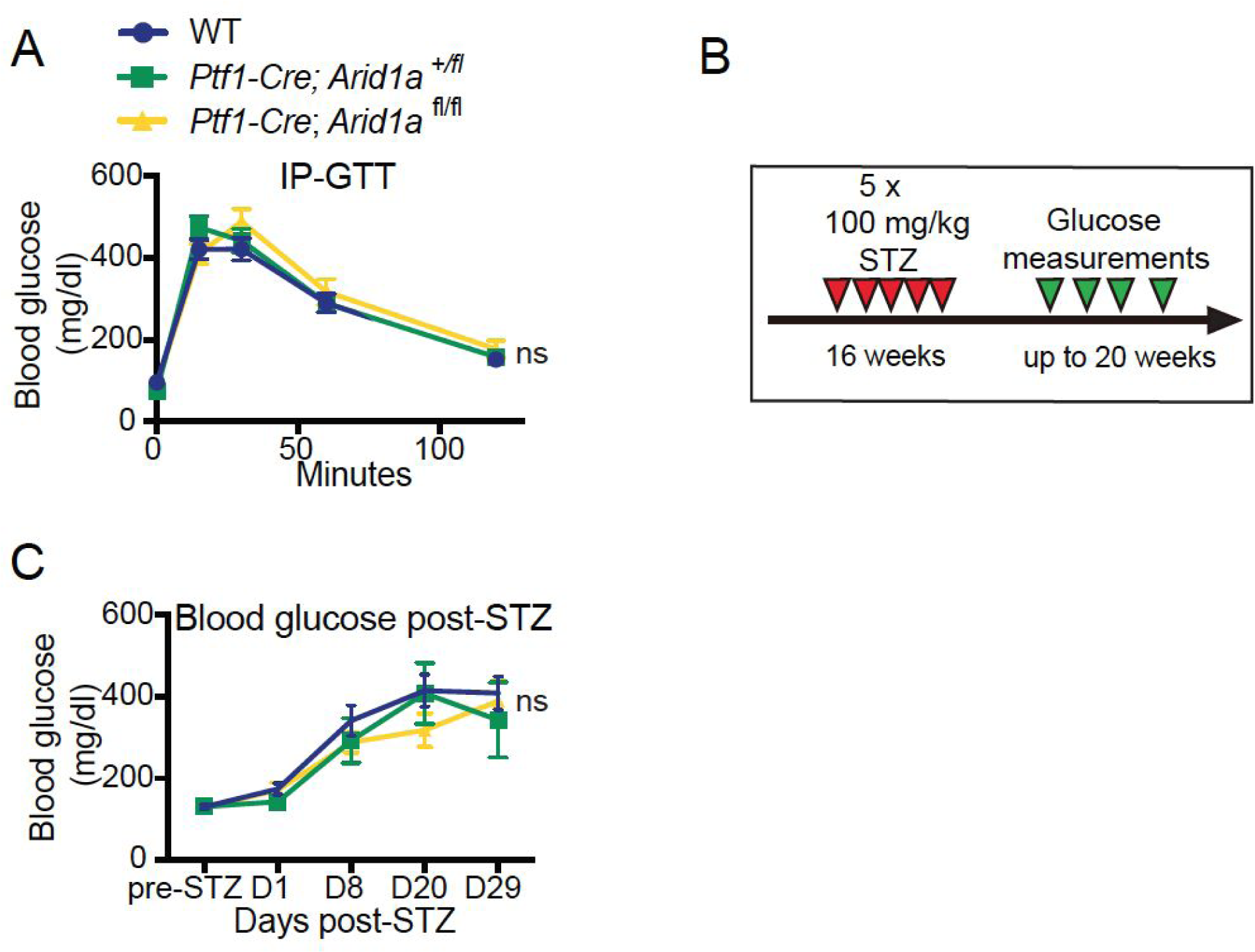
Acinar and ductal knockout of *Arid1a* does not phenocopy the whole body knockout. **A.** Intraperitoneal glucose tolerance test for *Arid1a^Fl/Fl^, Ptf1a-Cre; Arid1a^+/Fl^, and Ptf1a-Cre; Arid1a^Fl/Fl^* mice after overnight fasting. **B.** Experimental timeline for STZ-induced diabetes. **C.** Fed state blood glucose measurements taken post-STZ.

**Figure S2.**
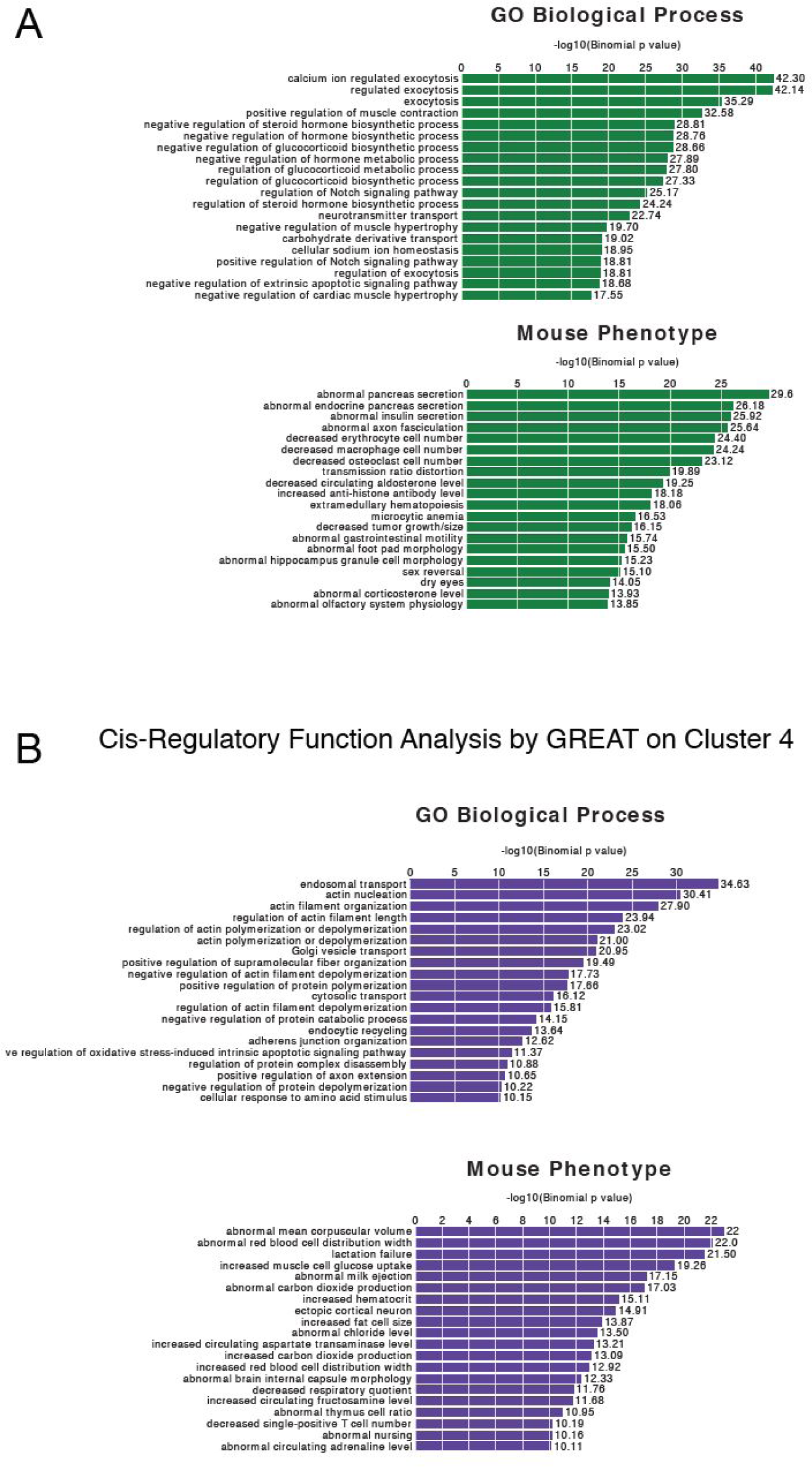
Cis-regulatory function analysis by GREAT for H3K27Ac peaks. **A.** Cis-regulatory function analysis by GREAT on sites within Cluster 3 (increased H3K27Ac in *Arid1a* HET condition) given as GO processes and mouse phenotypes. **B.** Cis-regulatory function analysis by GREAT on sites within Cluster 4 (increased H3K27Ac in *Arid1a* KO condition) given as GO processes and mouse phenotypes.

**Figure S3.**
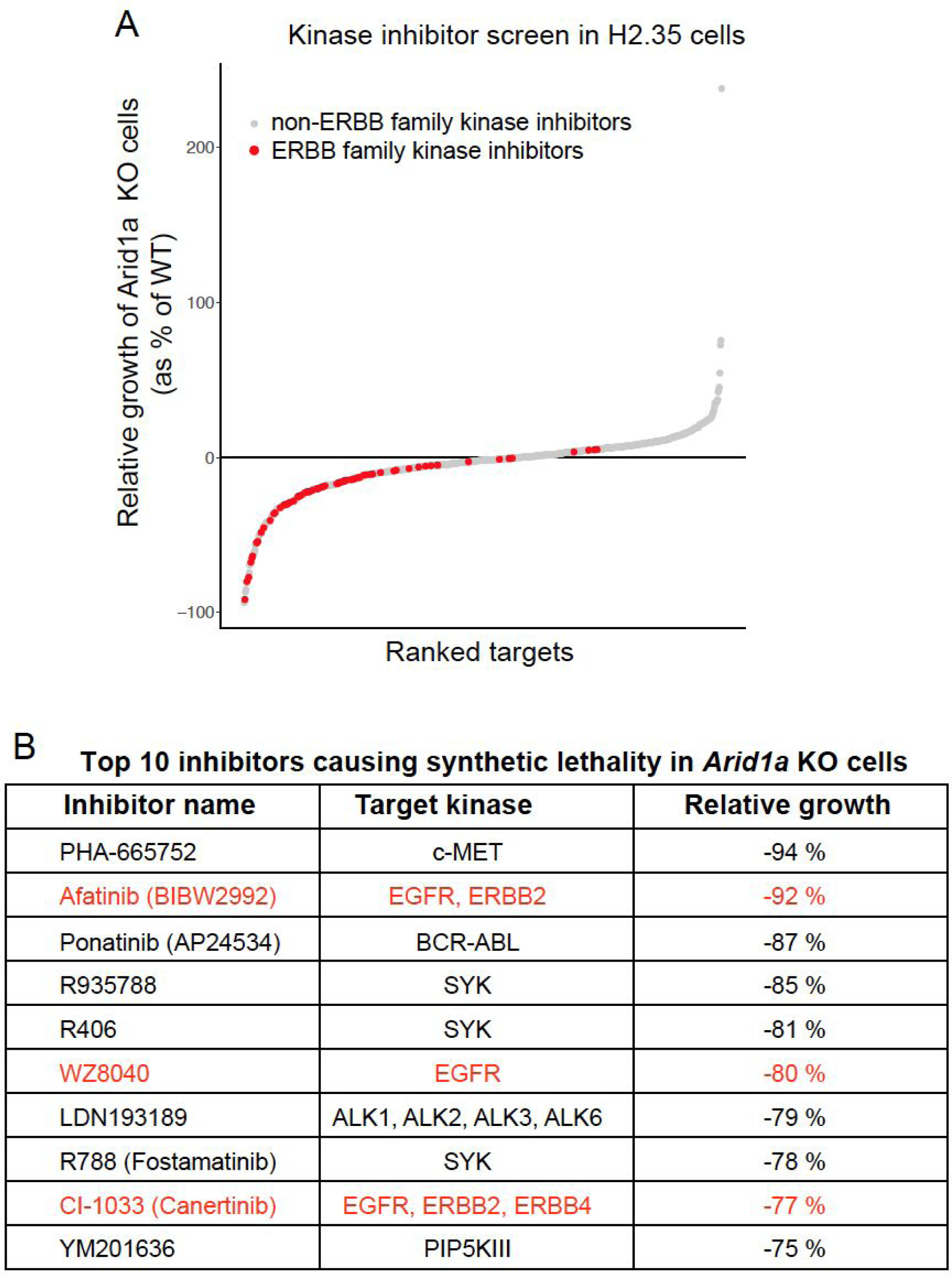
Kinase inhibitor screen in H2.35 cells shows that *Arid1a* deficient cells are more sensitive to cell growth inhibition induced by ERBB inhibitors than WT cells. **A.** Relative cell growth (KO vs WT). About 300 kinase inhibitors were used in this screen. Each gray dot corresponds to relative growth of *Arid1a* KO vs. WT cells in the presence of a non-ERBB kinase inhibitor, whereas each red dot corresponds to an ERBB family kinase inhibitor. **B.** Table showing the top 10 kinase inhibitors that caused the highest reduction of growth/viability selectively in *Arid1a* KO cells. 3 out of 10 top kinase inhibitors targeted the ERBB family of RTKs.

